# Impact of serum versus anticoagulant-containing plasma on influenza virus neuraminidase-based serological assays

**DOI:** 10.1101/2025.06.07.658460

**Authors:** Thi Hoai Thu Do, Stephen J. Kent, Adam K. Whealtey, Marios Koutsakos

## Abstract

The influenza virus neuraminidase (NA) is a promising target for next-generation influenza vaccines but standardized protocols for NA-based serological assays are lacking. Previous studies have demonstrated discordant results from haemagglutination inhibition and live virus microneutralization assays when comparing matched serum and plasma samples. It is therefore important to consider is the choice of serum or plasma samples in assays measuring influenza virus NA-specific antibodies. Here, we compared antibody titres against influenza A and B virus NAs in matched serum and different types of plasma using an enzyme-linked lectin assay (ELLA) and an enzyme-linked immunosorbent assay (ELISA). We observed good correlations between titres determined in serum and different types of plasma. However, there was variable and often poor agreement in the nominal titre values obtained from serum and different kinds of plasma in both ELLA and ELISA, with plasma samples often resulting in lower titres compared to serum samples. We also found differences in NA-specific responses to seasonal influenza vaccination assessed in serum versus plasma. Overall, our data demonstrate discrepancies between NA-specific antibody measurements in serum and plasma. We recommend the consistent use of serum as an important part of standardising NA-based serological assays.

## Introduction

Current influenza vaccines are formulated to target the hemagglutinin (HA) proteins of both influenza A and influenza B viruses. Vaccine strains are selected based on influenza surveillance and antigenic characterisation of circulating viruses. However, vaccine effectiveness can be limited by mismatch of circulating strains and those included in the vaccine [1-3]. Inclusion of the influenza virus neuraminidase (NA) in the vaccine has been suggested due to the associations of NA-specific antibodies and protection from influenza viruses and to increase the breadth of protection in years of HA mismatch [4, 5]. Indeed, influenza vaccines containing NA are being tested in human clinical trials [6-8].

Standardisation of serological assays to measure antibodies against the influenza NA is an important aspect for the development of vaccines that include NA. For the measurement of anti-NA antibodies, two main approaches are available: an enzyme-linked immunosorbent assay (ELISA) to determine binding antibodies, and an enzyme-linked lectin assay (ELLA), to measure inhibition of NA activity. With NA-containing vaccines in clinical trials, there is increased need for standardisation of serological assays to measure NA-specific immunity, with some protocols already developed [9, 10]. As part of this standardisation, it is important to consider the impact of the specimen type used. While serum is frequently used for most of assays measuring antibodies, plasma maybe sometimes preferable due to constrains on blood volume collection in human studies and the preference of plasma for measurement of various other analytes in blood. In addition, plasma may be available from historical cohorts that may warrant retrospective analysis of NA-specific antibodies. However, anticoagulants present in plasma may interfere with the readouts of serological assays [11, 12]. Indeed, specimen type has been shown to impact the determination of hemagglutination inhibition (HAI) and microneutralization (MNT) titres against influenza viruses [11, 12].

Given the effect of specimen type on NA inhibition (NAI) titres has not been investigated, we compared influenza NA-specific antibodies between serum and different types of anticoagulant-containing plasma in ELISA and ELLA.

## Materials and methods

### Viruses & recombinant NA proteins

A/California/07/2009 H1N1 viruses were propagated in 10-day old embryonated chicken eggs. B/Malaysia/2506/2004 viruses were propagated in MDCK cells. Recombinant NA proteins consisting of the ectodomain (head domain) of NA linked with a hexa-Histidine affinity tag (HisTag) and an AviTag were produced in house as previously described [13, 14].

### Human serum samples and plasma samples

Samples were collected under study protocols that were approved by the Alfred Health Ethics Committee (ID 432/14) and the University of Melbourne Human Research Ethics Committee (ID 1443420). All participants provided written informed consent in accordance with the Declaration of Helsinki. For the 2015, 2016 and 2022 cohorts, serum and heparinised blood were collected prior to and 28 days post-vaccination. In 2024 cohort, blood samples from sixteen healthy donors were collected using 4 different blood collection tube including serum tube clot activator (Greiner, REF454071), K3-EDTA tube (Greiner, REF454036), 3.2% sodium citrate tube (Greiner, REF454327), and lithium heparin tube (Greiner, REF454084). Sera and plasma were isolated and cryopreserved at -80°C.

### ELLA for NAI titre determination

ELLA was performed as previously described [15]. Firstly, 100μl of of 25μg/ml Fetuin from fetal bovine serum (Sigma) were coated on Maxisorp 96-well plates (Thermo Fisher) at 4°C overnight. For the measurement of NA activity, viruses were serially diluted 2-fold in assay buffer (33 μM MES, 4 mM CaCl_2_, pH 6.5, 1% BSA, 0.5% Triton X-100) starting from 1:2 dilution. Next, 25μl of assay buffer followed by 25μl of diluted virus were added into Fetuin-coated Maxisorp plate and incubated at 37°C for 16-18 hours. Then, 100μl of horseradish peroxidase-conjugated arachis hypogaea lectin (PNA-HRP) (Eylabs) was added at concentration of 1μg/ml and incubated for 2 hours at room temperature. Between each step, plates were washed 6 times with PBS-T (PBS with 0.05% Tween-20). Plates were developed using 100μl TMB substrate (Sigma) for 3 minutes, stopped by 50μl 0.16M sulphuric acid and read at 450 nm. Plates were read at OD450nm. The EC_70_ value of viruses were determined by fitting a curve with non-linear regression in Prism 9 (GraphPad)and identifying the value corresponding to 70% of NA activity. These values were used for NA inhibition assay.

For the determination of NA inhibition activity of serum and plasma, viruses were diluted based on EC_70_ value, as determined above. Serum and plasma samples were treated with RDE and heat-inactivated for 1 hour. For A/California/07/2009 H1N1 viruses, serum and plasma samples were serially diluted 2-fold starting from 1:10 dilution. For B/Malaysia/2506/2004 viruses, serum and plasma samples were serially diluted 3-fold starting from 1:40 dilution. Then, 25μl of diluted sera or plasma was added into Fetuin-coated Maxisorp plates, following by an addition of 25μl of diluted viruses. The mixture was incubated at 37°C for 16-18 hours. Wells with no viruses were used as negative controls. Wells with no serum were used as positive controls. Plates were developed using 100μl TMB substrate (Sigma) for 3 minutes, stopped by 50μl 0.16M sulphuric acid and read at 450nm. An IC_50_ value was calculated by interpolating the value of serum/plasma dilution that correspond to NAI of 50% using Prism 9 (GraphPad). An HA stem-specific antibody (CR9114) was used to confirm the absence of HA-specific antibody interference in the ELLA.

### ELISA

ELLA was performed as previously described [15]. Recombinant NA proteins were coated on 96-well Maxisorp plates (Thermo Fisher) at concentration of 2 μg/ml overnight at 4°C. Plates were firstly blocked with 200μl of 1% FCS in phosphate-buffered saline (PBS). Sera and plasma samples were serially diluted 4-fold starting from 1:100 dilution and a 100μl was added to each well for 2 hours at room temperature. Then, 100μl of HRP-conjugated rabbit anti-human IgG (1:20,000; Dako), was added for 1 hour at room temperature. Between each step, plates were washed in PBS-T (0.05% Tween-20 in PBS) and PBS. Plates were developed using 100μl TMB substrate (Sigma) for 3 minutes, stopped by 50μl 0.16M sulphuric acid and read at 450 nm. The graph of OD450nm corresponding to dilution of serum/plasma samples was determined by fitting a curve with non-linear regression in GraphPad Prism. Endpoint titres were calculated as the reciprocal serum dilution giving signal 2x background using a fitted curve (4 parameter log regression) in Prism 9 (GraphPad).

### Statistical analysis

Log_10_-transformed titres were used for analysis. Data were tested for normality using the Anderson-Darling test and correlation was assessed by computing nonparametric Spearman correlation. Assay agreement was determined using Bland-Altman analysis. Significance of differences between groups was assessed by a one-way ANOVA test and Dunnett’s multiple comparisons tests or a two-way ANOVA test and Sidak’s multiple comparisons tests. All statistical analyses were performed in Prism 9 (GraphPad).

## Results

### Determining the level of acceptable agreement between sample types

To determine if serum and plasma samples can be used interchangeably, we firstly needed to determine an acceptable level of agreement between the two sample types. We considered a two-fold difference two be acceptable as this is a commonly used threshold in serological assays [11, 12, 16, 17]. This would only be appropriate if the assays tested in our study show good repeatability with less the 2-fold variation between repeat measurements. Therefore, we firstly assessed the repeatability of the ELLA assay for serum and plasma by performing independent repeat measurements of NAI titres using 28 serum and 28 plasma samples obtained pre- and post-seasonal influenza vaccination in 2016 from 14 individuals. A B/Malaysia/2506/2004 recombinant NA (rNA) protein was used based on availability. NAI titres were positively correlated between two independent experiments for both serum (r=0.9) and plasma (r=0.89) **(Figure 1A)**. While the two measurements show good correlation, this does not mean the two measurements provide the same nominal titre values, which would make them interchangeable. We therefore used the Bland-Altman analysis for agreement between measurements **(Figure 1B)**, commonly used to determine agreement between serological assays [11, 12, 18, 19]. The Bland-Altman analysis produces a graph with the average of the two readings plotted against the difference of the same two readings (referred to as bias). A bias value close to zero indicates good agreement. Using Bland-Altman analysis, we found that the average bias in log_10_IC50 between repeated measurements was 0.05 for serum (corresponding to 0.9-fold difference) and -0.08 for plasma (corresponding to 1.19-fold change). We found that 96% of measurements in serum and 85% of measurements in plasma were less than 2-fold different between the two experiments. Therefore, in subsequent analyses comparing serum and plasma, we considered a difference of ±0.3 in log_10_ titres (corresponding to a 2-fold difference) as indicative of acceptable agreement between sample types.

**Figure 1:**
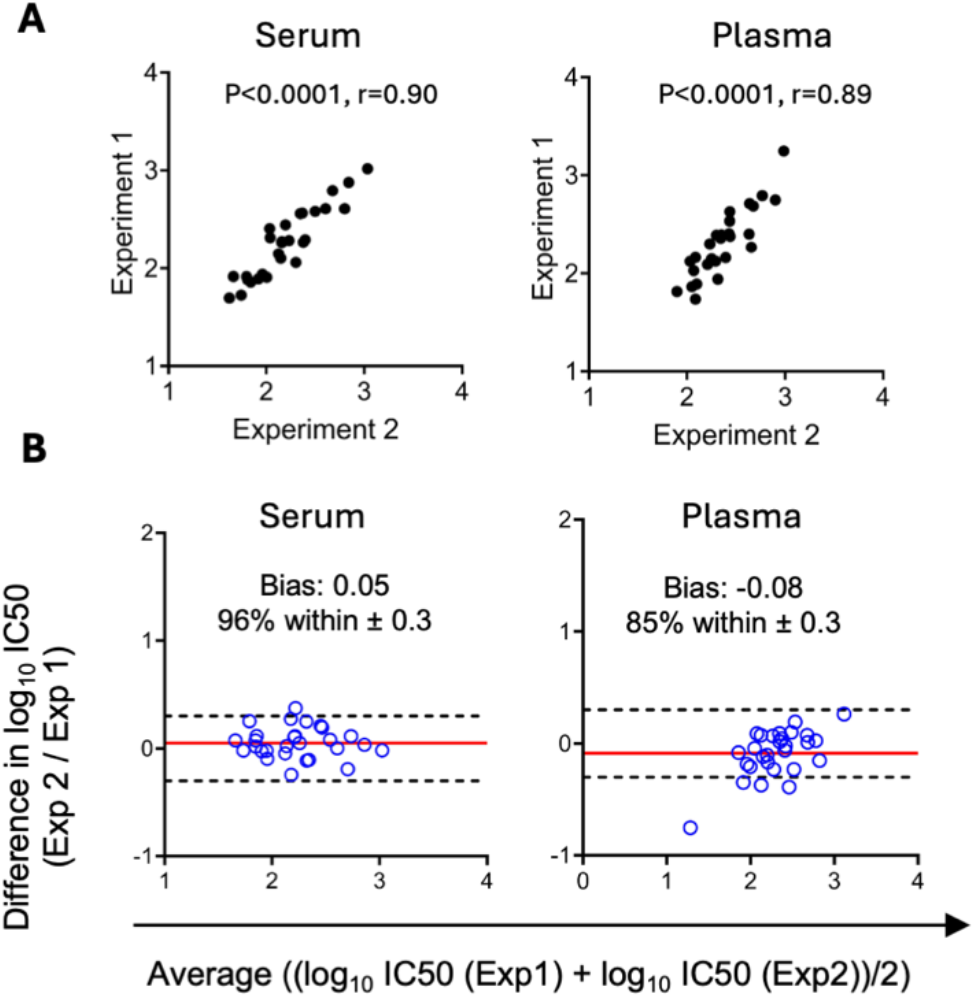
The NAI activities of heparin-plasma and serum against B/Malaysia/2506/2004 from vaccinated donors were measured by ELLA at baseline and 28 days post-vaccination. Donors from 2016 cohort (n=14) were tested. Each sample was tested in 2 independent experiments. **(A)** Spearman correlation in NAI titres between 2 experiments was assessed. **(B)** Agreement in NAI titres between 2 experiments was assessed by Bland-Altman analysis. The difference between titre values obtained from 2 experiments is plotted on the y-axis and the average of the two values is plotted on the x-axis. The red lines indicate the mean difference between two values (bias) and dash lines indicate the limit of agreement within ± 0.3 (log_10_ 2-fold different).

### Variable agreement in NAI titres and vaccine responses to NA measured in serum and plasma

We started with evaluating the level of agreement between titres in matched serum and heparin-containing plasma in baseline and post-vaccination samples from 3 cohorts of seasonal influenza vaccination (n=22 from 2015; n=14 from 2016, n=11 from 2022). **(Figure 2A-B)**. Using Bland-Altman analysis of NAI titres, a bias value close to zero indicates good agreement and we considered a bias value ± 0.3 as an acceptable level of agreement as determined above. A positive bias value indicates lower NAI titres in plasma compared to serum and negative bias value indicate higher NAI titres in plasma compared to serum. Using Bland-Altman analysis, we observed variable agreement between serum and plasma across cohorts and pre- or post-vaccination samples with 3 instances of a mean bias outside of the ± 0.3 cut-off. In addition to the average bias, we also considered the percentage of average difference values that lied outside of the ± 0.3 cut-off. This ranged from 14-54% across cohorts and pre- or post-vaccination samples **(Figure 2A)**. Therefore, across all 6 comparisons, less than 86% of matched serum and plasma samples were within the acceptable range of agreement. Additionally, we observed significantly lower NAI titres in plasma compared to serum in the 2015 cohort at baseline (serum with mean log_10_ NAI = 1.77, plasma with mean log_10_ NAI = 2.18) and in the 2022 cohort post-vaccination (serum with mean log_10_ NAI = 2.85, plasma with mean log_10_ NAI = 2.62). Meanwhile, significantly higher NAI titres were observed in plasma compared to serum at baseline of cohort 2022 (serum with mean log_10_ NAI = 2.55, plasma with mean log_10_ NAI = 2.72) **(Figure 2B)**. Overall, we found variable and often limited agreement in NAI titres determined in matched serum versus plasma.

**Figure 2:**
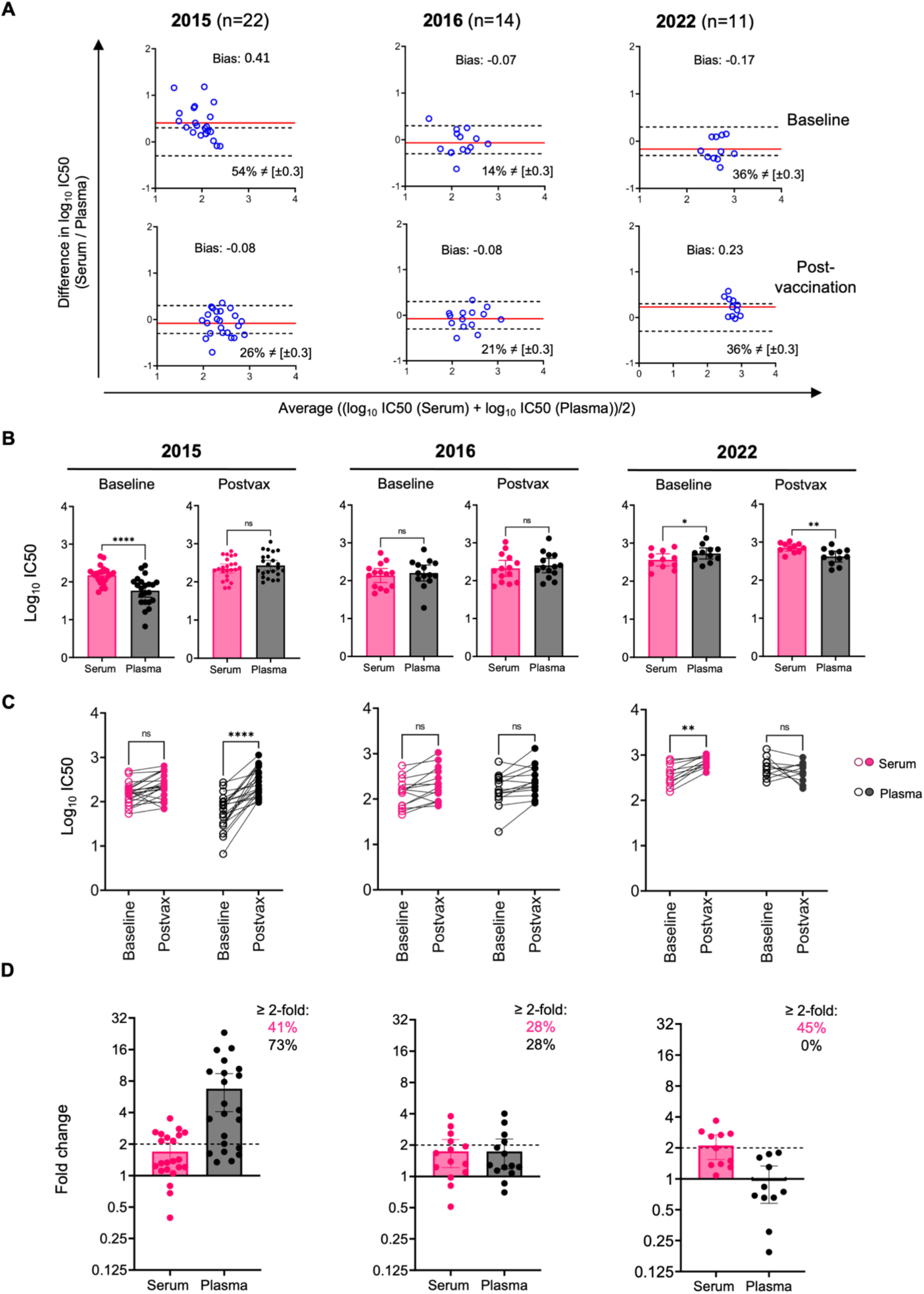
The NAI activities of heparin-plasma and serum against B/Malaysia/2506/2004 from vaccinated donors were measured by ELLA at baseline and 28 days post-vaccination. Donors from 3 cohorts were tested: 2015 (n=23), 2016 (n=14), 2022 (n=11). **(A)** Bland-Altman plots comparing NAI titres between serum and plasma samples was shown. The difference between titre values obtained from plasma and serum is plotted on the y-axis and the average of the two values is plotted on the x-axis. Dash lines indicate the limit of agreement withing ± 0.3 (log_10_2). The red line indicates an average difference between serum versus plasma (bias). Positive bias suggests overall higher titres and negative bias suggests overall lower titres of serum compared to plasma. **(B)** NAI titres of serum and plasma pre- and post-vaccination were plotted. The significance of differences between groups of samples was assessed by paired t-test. **(C)** NAI titres of serum and plasma pre- and post-vaccination were plotted. The significance of differences between groups of samples was assessed by two-way ANOVA test and Sidak’s multiple comparisons tests. **(D)** Fold-change in NA responses at post-vaccination compared to baseline were shown.

Next, we investigated the extent to which the variability between serum versus plasma affect the evaluation of vaccine responses **(Figure 2C)**. NAI titres to B/Malaysia/2506/2004 increased from pre-to post-vaccination in plasma but not serum samples in the 2015 cohort, and in serum but not plasma samples in the 2022 cohort. No significant changes were observed after vaccination in either serum or plasma samples in the 2016 cohort. The distribution of NAI responses (fold-change in titre relative to pre-vaccination) in plasma and serum samples (**Figure 2D**) varied between the two sample types in the 2015 and 2022 cohorts but not in the 2016 cohort. We observed that the mean fold-change measured in plasma was different from the mean fold change in serum samples in the 2015 and 2022 cohorts. The mean fold-change measured in plasma was also highly variable across the three cohorts (6.75-fold for 2015, 1.74-fold for 2016, 0.96-fold for 2022), while the NAI response determined in serum was more consistent across three cohorts (1.69-fold for 2015, 1.74-fold for 2016, 2.10-fold for 2022). Additionally, the percentage of individuals with a 2-fold or higher increase in NAI also varied between the two serum and plasma (41% versus 73% in 2015; 28% versus 28% in 2016; 45% versus 0% in 2022). Overall, we observed variability in NAI vaccine responses between matched serum and plasma samples.

In summary, these data demonstrate that the level of agreement NAI titres and vaccine responses vary considerably between serum and heparin-containing plasma.

### Limited agreement of NAI titres and NA endpoint titres to IAV N1 and IBV NA between serum and different types of plasma

We next extended our analysis to include other types of possible anticoagulants like 3.2% sodium citrate and K3 EDTA collected from 16 donors in 2024 prior to any vaccination, in addition to serum and heparin plasma. We also evaluated two different influenza strains (A/California/07/2009 H1N1 and B/Malaysia/2506/2004) and assessed both NAI by ELLA and NA binding titres using ELISA. We found significantly lower ELISA titres to N1 in EDTA samples (mean log_10_titre = 2.98) compared to serum (mean log_10_titre 3.4), and significantly lower NAI titres to IBV NA in EDTA (mean log_10_titre = 2.48) and sodium citrate (mean log_10_titre = 2.75) samples compared to serum (mean log_10_titre = 3.03) **(Figure 3A)**. In both ELISA and ELLA, antibody titres were positively correlated between serum and different types of plasma, as well as between the different types of plasma (r >0.7, p<0.001) **(Figure 3B-C)**. Using Bland-Altman analysis of ELISA titres **(Figure 4A-B)**, we found poor average agreement (bias beyond ±0.3) only for N1 between Serum and EDTA plasma. However, across all comparisons of N1-specific titres 19-75% of average difference values were greater than 2-fold. ELISA titres to BNA showed better agreement, with average biased within the ±0.3 threshold and on 6-19% of average difference values being greater than 2-fold. Agreement in ELLA titres was poor for all comparisons, with 19-81% of average difference values being greater than 2-fold. Overall, although NAI and ELISA titres are positively correlated between serum and plasma, our data demonstrated variable and often limited agreement between serum and different types of anticoagulant-containing plasma across 2 influenza NA antigens.

**Figure 3:**
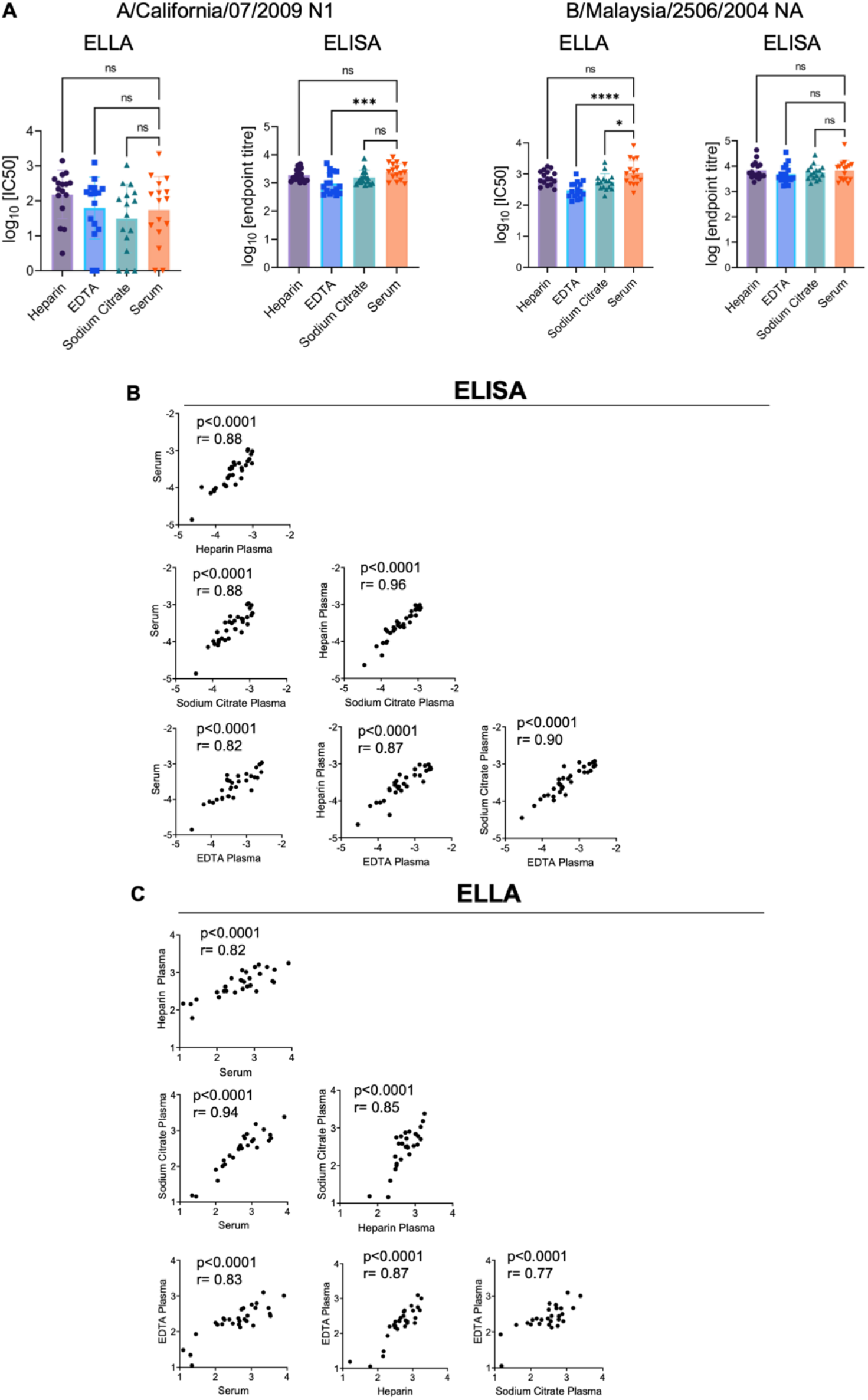
The NAI activities of EDTA plasma, heparin plasma, sodium citrate plasma, and serum from 16 donors were measured by ELLA and ELISA. For ELLA, two virus strains were tested: A/California/07/2009 H1N1 and B/Malaysia/2506/2004. For ELISA, two rNA proteins were tested: A/California/07/2009 N1 and B/Malaysia/2506/2004 BNA. **(A)** The summary of NAI titres from ELLA and NA endpoint titres from ELISA of EDTA plasma, heparin plasma, sodium citrate plasma, and serum from 16 donors was shown. The significance of differences between groups of samples was assessed by one-way ANOVA test and Dunnett’s multiple comparisons tests. **(B-C)** Correlations between serum and different types of plasma in ELISA and ELLA. The Spearman correlation between each serum/plasma pair was assessed (n=32, from 16 donors and 2 tested virus strains/rNA proteins).

**Figure 4:**
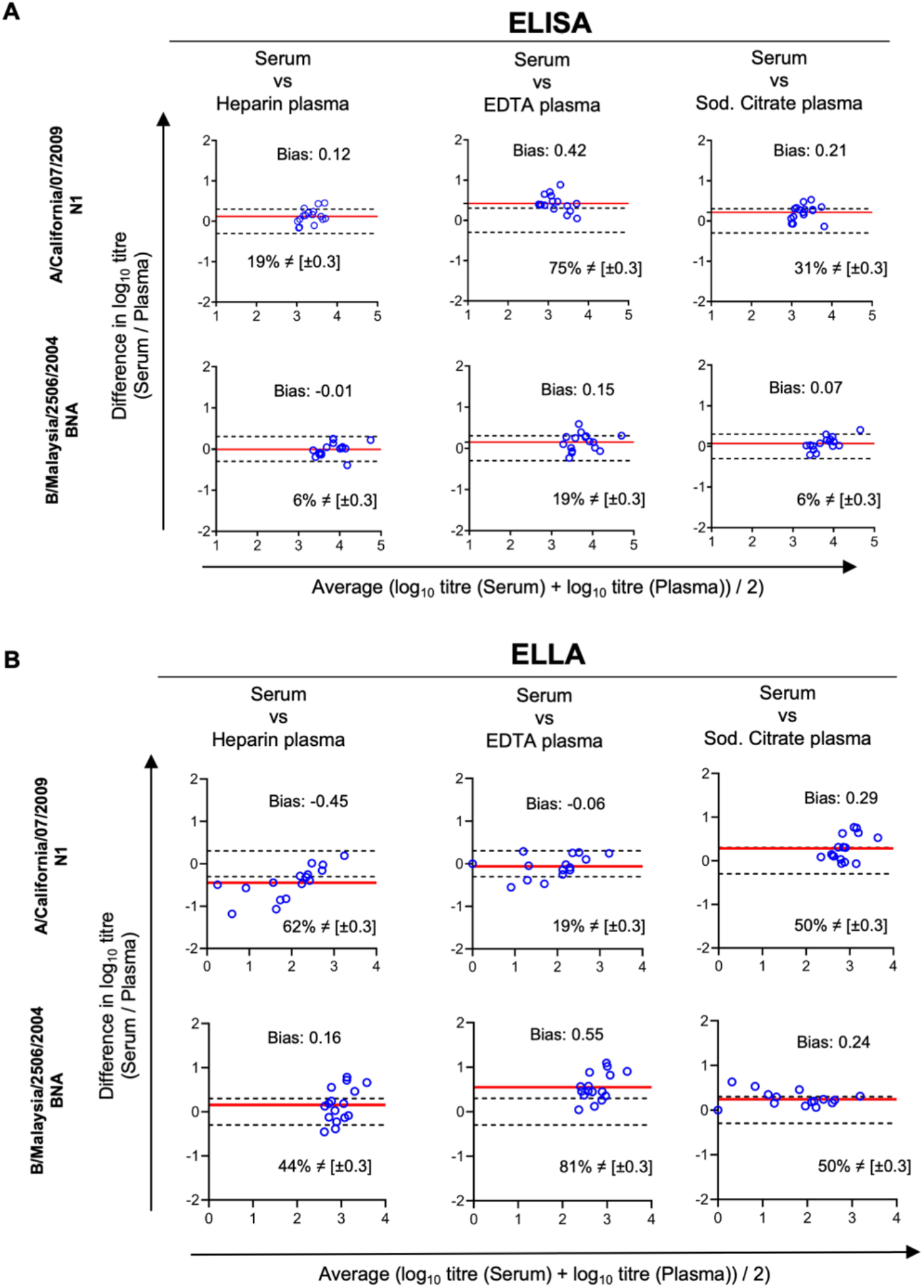
**(A)** Bland-Altman plots comparing NAI titres in ELISA for each type of plasma compared to serum. **(B)** Bland-Altman plots comparing NA endpoint titres in ELLA for each type of plasma compared to serum. The difference between titre values obtained from plasma and serum is plotted on the y-axis and the average of the two values is plotted on the x-axis. Dash lines indicate the limit of agreement within ± 0.3 (log_10_2). The red line indicates an average difference between serum versus plasma (bias). Positive bias suggests overall higher titres and negative bias suggests overall lower titres of serum compared to plasma.

## Discussion

Standardised serological assays for determining NA-specific antibodies will be key in characterizing the immunogenicity of next-generation influenza vaccines incorporating NA. An important factor to consider is the type of sample used (serum versus plasma). For serum and plasma samples to be used interchangeable they should provide acceptably similar titre values. Throughout our analysis with different types of plasma from different cohorts, different antigens (N1 and BNA) and different assays (ELLA and ELISA) we found poor agreement in the titres determined in serum and matched plasma, despite titres being generally well correlated between sample types. Overall, these data demonstrate that the anticoagulants present in plasma may affect the measurement of NA-specific antibodies and suggest the consistent use of serum across studies.

Our study clearly demonstrates that while NAI titres measured in plasma and serum are well correlated, the different sample types provide different nominal values. Indeed, across all the comparisons of NAI titres across antigens, sample types and cohorts **(Figures 2A and 4B)**, less than 86% of average difference values varied by less than 2-fold. In addition to different nominal titre values, NAI vaccine responses were also different when determined using serum and heparin containing plasma. While the influenza vaccine is not standardised for NA content, residual NA can stimulate NA-specific antibodies [20, 21]. NA-specific antibody responses (measured by ELLA) varied between serum and heparin plasma in 2/3 cohorts tested. We note that the direction of agreement varied depending on the year with greater NA responses in plasma than serum in 2015, greater NA responses in serum than plasma in 2022 and no difference in 2016. This variability between years may relate to storage of samples across different years or batch effects of blood collection tube. We also note, however, that NA responses to vaccination were consistent in serum (fold-change ranged from 1.69-2.10 with 28-45% of individuals with >2-fold increase across the 3 years) but highly variable in plasma samples (fold-change ranged from 0.96 to 6.75 with 0-73% of individuals with >2-fold increase across the 3 years). This may result from the differences in baseline titres observed in plasma **(Figure 3A-C)**. Overall, NAI responses to seasonal influenza vaccination differed between serum and heparin plasma, further supporting the preferential and consistent use of serum in influenza vaccine immunogenicity trials.

While we are not aware of previous studies comparing serum and plasma in NA serological assays, others have compared HAI or MN titres measured in serum and plasma and found variable effects across EDTA plasma, heparin plasma and sodium citrate plasma compared to serum [11, 12]. EDTA plasma was shown to cause overestimation of titres in both HAI and MNT assays [11], while sodium citrate plasma and heparin plasma were demonstrated to cause underestimation of HAI titres [12]. Our results with NAI titres further demonstrate the variable impact of anticoagulants on serological assays. The differences in titres determined in serum and plasma may relate to the anti-coagulants used in plasma collection. EDTA and sodium citrate can act as chelating agents [11, 22, 23] and may limit ions important for assay activity. Heparin has been previously reported to interfere with the interactions between antibodies and antigens [11, 24].

A limitation of our study is the limited number of samples tested for each comparison (n=11-22). Nonetheless, across the different cohorts and assays tested, we consistently observed moderate to poor agreement between serum and plasma. In addition, our analyses were conducted primarily using the B/Malaysia/2506/2004 NA antigen, selected based on availability. However, our analyses using the N1 antigen from IAV further confirmed the limited agreement between serum and plasma.

In conclusion, while both serum and plasma can be used to measure NA-specific antibodies, caution is warranted in the use of plasma samples as they show variable and often limited agreement with matched serum samples. Importantly, when using titres from plasma or serum we observed differences in vaccine responses to NA. This might lead to an overestimation or underestimation of vaccine responses impacting the outcomes of immunogenicity trials. Given the impact of anticoagulants on NA serological assays, we recommend the consistent use of serum for NA serology and the consideration of sample type in the standardisation of NA serological assays.

## Acknowledgements

We thank the participants for their generous involvement and provision of samples. The work has been generously supported by the Morningside Foundation and by the Australian National Health and Medical Research Council. For the purposes of open access, the author has applied a CC BY public copyright licence to any Author Accepted Manuscript version arising from this submission.

## Author contributions

M.K and T.H.T.D designed the study. T.H.T.D performed experiments. T.H.T.D and M.K analysed data. A.K.W and S.J.K provided samples critical for the study. T.H.T.D, A.K.W, S.J.K, and M.K drafted the manuscript. All authors reviewed the final version of the manuscript.

## Competing interest

M.K. has acted as a consultant for Sanofi group of companies. The other authors declare no competing interests.

